# From periodic/aperiodic to static/dynamic: rethinking the decomposition of neural power spectra

**DOI:** 10.64898/2026.07.26.740811

**Authors:** Daniele Marinazzo

## Abstract

The power spectrum of electrophysiological signals exhibits both narrowband peaks and a broadband 1/f-like slope. The two are routinely treated as separable components: a "periodic" foreground and an "aperiodic" background; and each has been related to behaviour, cognition, and clinical status. This paper argues, and illustrates with simulations and case studies, that the boundary between the two is less sharp than the standard decomposition assumes: non-stationary amplitude modulation of an oscillation can inflate the apparent spectral trend (whereas clean periodic modulation does not, its sidebands being absorbed into the peak), and a broadband process viewed through any narrow band produces an envelope that mimics amplitude modulation. The two phenomena are mirror images of the same underlying property, that the temporal envelope and the spectral slope are coupled. I show what this means in practice with a set of illustrative scenarios and three case studies (a simulated alpha lateralisation paradigm, an auditory oddball ERP, and simultaneous scalp EEG and ECoG under rest and propofol sedation). I argue that the question "is the observed change a change in the aperiodic component" cannot in general be answered from time-series analysis alone, and that the temporal envelope of detected oscillations carries information that the static power spectrum loses. I ultimately argue that the conventional periodic/aperiodic partition of neural power spectra conflates a more fundamental distinction: between processes whose amplitude is stationary over the analysis window and processes whose amplitude changes. Transferring the dichotomy from the spectral domain to the time domain, from periodic/aperiodic to static/dynamic, reveals structure that the power spectrum alone cannot recover.

## 1. Introduction

The analysis of electrophysiological signals in the frequency domain has been one of the most productive frameworks in cognitive and clinical neuroscience. Narrowband rhythmic activity has been linked to a wide range of psychological processes and clinical conditions. These associations rest on the implicit assumption that the spectrum can be meaningfully divided into frequency-specific components.

The power spectrum of neural signals also exhibits a characteristic broadband decrease of power with increasing frequency: a spectral trend that is commonly fitted as a 1/f power law and labelled the "aperiodic" component, though, arguably, neither the power-law form nor its status as a separate component is secure. I will refer to it neutrally as the spectral trend, reserving "1/f" and "aperiodic exponent" for the specparam parametrisation and for others’ claims. Because the current literature overwhelmingly quantifies this trend with specparam, I nonetheless use the specparam exponent throughout as the operational measure of the trend, while treating the exponent as a descriptor of a decaying spectrum rather than as evidence of a genuine power law. This trend has two important practical implications. The first is methodological: the spectral trend acts as a frequency-dependent baseline, so any analysis comparing power across frequencies, or across conditions at a given frequency, risks conflating changes in the spectral trend with changes in narrowband oscillatory power. This motivated approaches that model the two components explicitly. The earliest of these, the Xi-Alpha model of Pascual-Marqui et al. (1988), is not merely a fitting convention but a generative proposal: the EEG spectrum is taken to arise from two physiologically distinct, additive processes: a broadband background (Xi), always present, and an intermittent oscillatory peak (Alpha), each estimated separately. Specparam (Donoghue et al., 2020) later operationalised the same periodic/aperiodic separation as a top-down parametric fit. The second is interpretive: the slope itself has been proposed as a functionally meaningful signal, linked to excitation-inhibition balance (Gao et al., 2017), membrane timescales (Gao et al., 2020), and arousal state, effectively treating the 1/f component as a distinct frequency band with its own physiological significance.

A complementary frequency-domain approach to the same separation problem is IRASA (Irregular-Resampling Auto-Spectral Analysis; Wen & Liu, 2016), which isolates the fractal from the oscillatory component by exploiting self-affinity: resampling a fractal process by a non-integer factor scales its power spectrum predictably, while oscillatory peaks shift in frequency, so the median across resampling factors recovers the broadband background. IRASA makes explicit the same power-law assumption that specparam makes implicitly, but reaches the separation by operating on the time series rather than fitting a parametric model to the spectrum; a detailed empirical and simulation-based comparison of the two methods, including shared failure modes such as overlapping peaks and low-frequency spectral plateaus, is given by Gerster et al. (2022). Neither specparam nor IRASA, however, takes the temporal structure of the amplitude explicitly into account: both summarise the whole recording as a single spectral decomposition, and none of these methods individually distinguishes a stationary broadband process from a non-stationary oscillatory amplitude as the source of an observed 1/f-like slope.

Moving to the detection of transient activity, the fBOSC method (Seymour et al., 2022) uses specparam-calibrated power thresholds to detect oscillatory bursts, providing burst abundance and duration statistics that carry information about the temporal structure of oscillatory activity; it builds on the extended BOSC family (eBOSC; Kosciessa et al., 2020), which introduced robust background estimation and the grouping of supra-threshold timepoints into rhythmic episodes with explicit duration tracking. The cycle-by-cycle approach of Cole & Voytek (2019) operates directly in the time domain, characterising individual oscillatory cycles by period, amplitude, and rise–decay asymmetry without reference to the power spectrum, and is therefore sensitive to waveform non-sinusoidality and within-cycle amplitude modulation that spectral methods cannot see.

Across these methods the convention is to treat the oscillatory peak and the aperiodic background as two separate quantities. Individual alpha frequency and alpha power describe the peak; offset and exponent describe the background; the two sets of parameters are reported side by side and related to behaviour, cognition or clinical status, often with the implicit or explicit assumption that they index independent neural mechanisms (Donoghue et al., 2020; Schaworonkow, 2023; Marsicano et al., 2026). The position taken in this paper departs from that convention: I treat at least some of what is reported as an independent change in the aperiodic component as, at the level of the observable signal, a re-expression of changes in oscillatory amplitude over time. The substantive critique of the "parallel mechanisms" reading is taken up in Section 4.5; the immediate concern of this paper is observational.

The emphasis on separating the "periodic" from the "aperiodic" has, however, introduced a conceptual sharpness that may not be warranted. Spectral parameterisation makes the standard stationarity assumption explicit and adds a further one: that the broadband slope and the oscillatory peaks are separable, additive, and independently interpretable. Yet the very act of parameterisation does not eliminate the underlying mixing. A signal whose oscillatory amplitude varies over time will, in a static power spectrum, produce a broadened peak and an inflated trend; these are not independent features of the data but different manifestations of the same underlying non-stationarity. The label "aperiodic" can give the false impression of always reflecting a distinct neural process, separate from oscillatory dynamics, up to the extreme of being called "brain dark matter" (Landau, 2021), when in fact it may simply reflect the temporal envelope structure of those very oscillations.

At the level of observable data, periodic and aperiodic components mix through amplitude modulation: any non-stationary oscillatory amplitude change produces spectral spreading (sidebands) into the spectral trend. The converse is equally true: a broadband 1/f process, when viewed through any narrowband filter, produces an amplitude envelope that fluctuates over time, mimicking an amplitude-modulated oscillation. The mixing is symmetric, and the periodic/aperiodic decomposition is not a separation of two distinct processes but a single signal viewed through two complementary lenses. At the level of mechanism, the two may also be continuous: cellular noise that drives oscillatory amplitude fluctuations is itself broadband, and what appears as a stationary broadband slope at the population level may be the incoherent superposition of many individually amplitude-modulated oscillatory processes at the cellular level. The question of which description is more appropriate depends on the scale of observation and the analysis window, not on a fixed property of the signal. That scale-free activity and rhythmic activity are entangled rather than independent is not a new observation: scale-free cortical activity carries nested cross-frequency structure (He et al., 2010; He, 2014), and 1/f-like spectra can be generated by superpositions of damped or relaxation processes (Muthukumaraswamy & Liley, 2018; Evertz et al., 2022; see Section 4.7).

A signal x(t) = A(t) cos(2π·f_0_·t + φ), where A(t) is a slowly-varying amplitude envelope, will produce a 1/f-like broadband slope in addition to a narrowband peak at f_0_. This is mathematically analogous in the power spectrum to a pure oscillation plus independent broadband noise; the two cannot be distinguished without examining the temporal structure of the amplitude envelope. The same ambiguity applies to oscillatory bursts, which produce spectral smearing around the carrier frequency that similarly inflates the apparent slope.

This degeneracy has a deeper and more practical dimension than is usually acknowledged. The spectral consequence of amplitude modulation depends critically on the nature of the modulator. A clean sinusoidal modulator at frequency f_m_ creates discrete sidebands at f_0_ ± f_m_, which specparam correctly absorbs into its peak model; in this case the aperiodic exponent is not inflated. But neural amplitude modulators are never clean sinusoids. In an ERP paradigm, oscillatory amplitude might ramp down after stimulus onset, stay suppressed, then recover; a non-stationary, ramp-like envelope change. In a task-versus-rest comparison, oscillatory amplitude shifts irregularly. Any such non-stationary amplitude change spreads energy continuously across a broad frequency range around the carrier, and specparam absorbs this spread into its aperiodic component, returning a positive exponent. This positive specparam exponent is not the same as a power-law (1/f) process: the magnitude depends on the envelope shape and is not generally a power law over the fitting range. This means that in practice, any study reporting an elevated aperiodic exponent in a task or ERP condition should consider whether the change reflects an alteration in broadband spectral structure, or simply an amplitude modulation of an oscillation whose envelope is non-stationary over the analysis window.

Oscillatory bursting is one instance of a more general phenomenon: any non-stationary amplitude modulation of an oscillation can change the specparam exponent in a way that can be mistaken for the signature of a broadband mechanism. Consider a signal consisting of a pure alpha oscillation interrupted by a beta burst: the alpha is suppressed during the burst, the beta appears transiently, and the resulting power spectrum shows an elevated aperiodic slope that is entirely a consequence of the non-stationarity of both oscillations’ amplitudes. No broadband source is required. A recent study has developed around detecting such bursts more accurately (Power et al., 2026), and it recognises that accurate burst detection requires first characterising the broadband background against which bursts are defined; methods such as PAPTO, eBOSC, fBOSC, and STPPTO all use specparam to estimate this background before thresholding. Power et al. (2026) also note that brief intermittent bursts, when trial-averaged, produce the appearance of sustained oscillatory power changes, a configuration in which the oscillatory envelope CV is exactly the diagnostic feature, regardless of whether the modulation is slow and smooth (classical ERD) or episodic and fast (burst on/off switching).

A key epistemic constraint shapes the analysis throughout this paper. The power spectrum, and any finite set of summary statistics derived from it, cannot uniquely identify the process that generated the signal. Multiple distinct generative processes can produce indistinguishable observable statistics. What can be assessed from the data is the temporal structure of the signal’s amplitude at the measurement scale: whether the oscillatory envelope is slow and modulated, episodic and bursty, or relatively stationary. These are observable descriptions, not claims about underlying circuit mechanism.

An important subtlety concerns the scale-dependence of stationarity. Whether a process appears stationary or amplitude-modulated depends on the length of the observation window relative to the timescale of the modulation. A slow AM process with a period much longer than the recording window will appear stationary within that window. At the extreme short end, individual action potentials are the canonical 1/f-spectrum signal when averaged across many cells and time, yet each spike is itself a deterministic event. The analyses presented here do not resolve this general and perhaps tautological question. They ask, instead: within a finite analysis window, does the temporal envelope structure of the signal show evidence of AM or bursting, or does it not? This is an operationally defined, scale-specific question that can be answered from the data.

A parallel, perhaps preliminary, perhaps more informative axis of description is then temporal: is the oscillatory amplitude stable over the analysis window, or does it fluctuate? Is the broadband slope consistent with a stationary stochastic process, or does it track amplitude modulation of a carrier? This reframing from periodic/aperiodic to static/dynamic motivates the analyses below.

The paper is organised as follows. Section 2 describes the data and the analysis methods, with the parameters of each method given alongside the data it was applied to. Section 3 illustrates how different temporal structures project into the same family of spectral statistics, using a small set of generative scenarios (Section 3.1–3.3) and three case studies: a simulated alpha lateralisation paradigm (Section 3.4), an auditory oddball ERP (Section 3.5), and simultaneous scalp and ECoG recordings under rest and propofol sedation (Section 3.6). Section 4 discusses implications, limits of the approach, and the open biophysical question of what the 1/f component actually reflects.

## 2. Methods

### 2.1 Data

#### Simulated scenarios

Five generative scenarios are used to illustrate how different temporal structures project into the same family of spectral statistics. None is intended as a model of any specific neural process or as a calibration target; the value is in showing what the features look like in each case.

All scenarios are 10 s sampled at f_s_ = 250 Hz. Signals were generated using the NeuroDSP toolbox (Cole et al., 2019) and combined by linear superposition. Pure oscillations: unit-amplitude sinusoids. Sinusoidal AM: carrier multiplied by (0.5 + 0.5 sin(2π × 1 Hz × t)), giving a positive envelope with mean 1 and modulation depth 50%. Broadband 1/f-like component: sim_powerlaw (exponent = −1), a stationary stochastic process with a power-law spectrum, used as a phenomenological broadband test case with no claim about biophysical mechanism. Beta bursts: sim_bursty_oscillation at 20 Hz with transition probabilities enter_burst = leave_burst = 0.1, yielding approximately 50% on-time. Additive Gaussian white noise (σ = 0.02) was also added to the time series.

The five scenarios are then: (A) pure alpha oscillation; (B) sinusoidally amplitude-modulated alpha; (C) alpha oscillation on a 1/f-like background; (D) bursty beta; (E) bursty beta on a 1/f-like background.

Additionally, an alpha lateralisation paradigm was simulated following Melcón et al. (2025). Two dipoles (contralateral and ipsilateral to a spatial cue) oscillated at a 10 Hz carrier across 60 trials, with epochs spanning −200 to 2000 ms relative to cue onset. After cue onset, the contralateral dipole underwent a Hann-shaped amplitude decrease (ERD: onset 350 ms, return 1,250 ms, depth 60%), while the ipsilateral dipole underwent a mirrored increase (ERS); a constant-amplitude dipole served as a control. Within each trial the carrier phase was drawn at random and shared across the three conditions, so that they differ only in their amplitude envelope. To isolate the effect of amplitude modulation on the spectral estimate, the signals were generated without additive noise or frequency jitter, and on a time base padded by 2 s on each side that was cropped to the −800 to 2000 ms display window after the wavelet transform, to remove edge artefacts. The variance-reduction rationale for trial averaging (Section 2.5) is general and does not depend on this noise-free construction.

#### Empirical datasets

These datasets are hosted on the NeuroTycho platform, www.neurotycho.org (platform paper Nagasaka et al. 2011); individual recordings are attributed to their original studies below (auditory oddball, Komatsu et al. 2015; resting scalp/ECoG, Oosugi et al. 2017).

Auditory oddball ERP: one channel on the auditory cortex of a marmoset (Fr), 80 trials (limited to the oddball ones), 256 Hz, −100 to +251 ms (Komatsu et al. 2015). The trial-averaged parameterisation was applied to the full trial matrix (5 ms step).

Resting-state recordings: simultaneous scalp EEG (channel T5) and ECoG (a channel from the posterior temporal area) recorded from a macaque monkey (Chibi), in two states (resting wakefulness and propofol sedation), 300 s at 256 Hz per condition (Oosugi et al. 2017).

### 2.2 Spectral parameterisation (specparam)

Power spectral density estimates were computed using Welch’s method with a window of 2 s for 10 s signals or 4 s for 300 s recordings (frequency resolution 0.25–0.5 Hz). For the auditory ERP grand average (0.35 s epoch), the full epoch was used as a single window. The fixed model (log P(f) = b − χ·log f) was used for all reported exponents over [1, 45] Hz. The knee model (log P(f) = b − log(κ + f^χ)) was also available but is not used for any reported result here; all reported exponents use the fixed model, and knee parameters and BIC comparisons are not reported.

The knee parameter has been interpreted as a membrane timescale (Gao et al., 2020), but it captures any spectral inflection in that range, including the inter-peak roll-off between two oscillations; it is not a direct read-out of membrane dynamics without corroborating evidence. For the resting-state recordings, specparam was applied both to the full 300 s spectrum (4 s Welch windows) and to 150 independent 2 s segments per condition to obtain the exponent distribution.

### 2.3 Time-resolved parameterisation (SPRiNT)

SPRiNT (Wilson et al., 2022) was used in MATLAB with fooof_mat as the specparam backend. The algorithm applies short-time Fourier transforms (1 s Hann window, 50% overlap) to a continuous recording, averages n_avg_ = 5 consecutive power spectra per time bin, and fits specparam at each bin. SPRiNT is applied here as a stationarity diagnostic: a stable exponent across bins is consistent with a stationary broadband process; a fluctuating exponent suggests non-stationarity due to AM or bursting.

### 2.4 Trial-averaged parameterisation

Assessing whether a change in the spectral trend reflects a change in broadband activity or a change in oscillatory amplitude over time is methodologically complicated in a short event-related epoch: SPRiNT’s short-time Fourier resolution (∼400–500 ms) is too coarse for effects unfolding over 100–300 ms, and applying specparam directly to a single-trial wavelet spectrogram requires heavy temporal smoothing to stabilise the per-bin fit, which trades away the resolution one is trying to gain. The approach used here instead applies specparam per time bin to a trial-averaged single-trial spectrogram, for event-related paradigms with epoch durations shorter than ∼4 s. For each trial, a complex Morlet wavelet convolution is performed at 1–44 Hz in 1 Hz steps, with n_cycles_ = 6, unit-norm normalised. The effective temporal resolution is set by the Gaussian envelope: FWHM(f) = n_cycles_ · √(2 ln 2) / (π · f). At 10 Hz this gives FWHM ≈ 225 ms; at 20 Hz, ≈112 ms; at 4 Hz, ≈562 ms. Instantaneous power is averaged across all N trials at each time-frequency point. By averaging across trials rather than across nearby time bins, the trial-averaged parameterisation reduces spectral noise by 1/√N without sacrificing temporal resolution; with N = 60 trials this gives a 7.7× noise reduction. Specparam (fixed mode, [1, 44] Hz) is fitted at every time bin (the timecourse is sampled every 10 ms for simulation, 5 ms for empirical data, which is the sampling step, not the resolution), yielding continuous timecourses of the aperiodic exponent, offset, R², and periodic peak parameters. The established approach to single-trial event-related parameterisation is to compute one power spectrum per trial over a fixed analysis window and fit specparam to each, then average the resulting parameters across trials; van Engen et al. (2026) use exactly this scheme, comparing two long windows (a pre-stimulus and a task window). This is well suited to slow, sustained effects, but a window long enough to yield a stable low-frequency spectral fit is much wider than a fast event-related transient, so any within-window change in the exponent is averaged out. The trial-averaged parameterisation used here differs only in the order of operations and in retaining time: instantaneous wavelet power is averaged across trials at each time-frequency point rather than each trial being reduced to a single windowed spectrum, and specparam is fitted per time bin rather than per window. This yields a continuous exponent timecourse at a resolution set by the wavelet width (∼225 ms at 10 Hz), which is what makes fast event-related changes such as the auditory N1, unfolding over tens of milliseconds, visible at all; it is otherwise the same specparam parameterisation applied to the same class of trial-based spectra. Averaging power spectra before fitting, rather than averaging the parameters of many noisier single-trial fits, also stabilises the estimate, since specparam parameters are non-linear functions of the spectrum and the mean of the fits need not equal the fit of the mean.

### 2.5 Amplitude envelope analysis

For each oscillatory band identified by specparam, the signal was bandpass-filtered (4th-order Butterworth) and the Hilbert analytic amplitude extracted, smoothed with a 3 Hz lowpass filter. The coefficient of variation CV = σ/μ of the smoothed envelope within the carrier band is sensitive to amplitude modulation: a sustained oscillation produces a near-flat envelope (low CV), an amplitude-modulated oscillation a slowly-varying envelope (higher CV). The CV must be computed within a specparam-detected carrier band, since bandpassing in an inactive band returns filter-induced fluctuations that do not reflect signal structure.

### 2.6 fBOSC burst detection

fBOSC (Seymour et al., 2022) uses specparam-calibrated power thresholds to detect time-frequency regions where power exceeds the expected level for a purely broadband process. In the taxonomy of Power et al. (2026), fBOSC belongs to the amplitude-based class alongside PAPTO, eBOSC, and STPPTO; all of which share the same prerequisite: an accurate estimate of the broadband background. Analysis frequencies: 4–44 Hz for simulated signals; 4–38 Hz for scalp EEG to exclude line noise. The χ²(2) threshold at the 95th percentile is pt(f) = χ²₀.₉₅,₂ · mp(f) / 2, where mp(f) is the aperiodic power. A contiguous epoch exceeding this threshold for at least 3 cycles is classified as a burst. Burst abundance (fraction of time in a burst at a given frequency) is the primary output used in this paper. When the broadband background is near zero, the threshold is trivially low and abundance approaches 1.0 for any oscillation; in this case fBOSC is not informative. For the resting-state recordings, fBOSC was computed on non-overlapping 5 s segments across the full recording, with burst abundance reported for the alpha (8–12 Hz) and beta (16–30 Hz) bands only.

## 3. Results

### 3.1 The spectral identifiability problem

Figure 1 illustrates the core problem. Two signals produced by different generative processes (an oscillation with non-stationary amplitude and no aperiodic component (panel A), and an oscillation superimposed on a stationary 1/f-like broadband process (panel B)) yield power spectra of similar overall shape. In both cases specparam summarises the spectrum as an alpha peak plus an aperiodic exponent. The exponents are not equal in magnitude, the broadband signal returns a higher value, but the decomposition assigns a positive exponent to the AM signal as well, and there is no feature of the spectrum or of the specparam output that identifies which generative process produced it. Two signals that differ structurally in time, one episodic, the other stationary, are summarised by the same family of spectral statistics.

**Figure 1.**
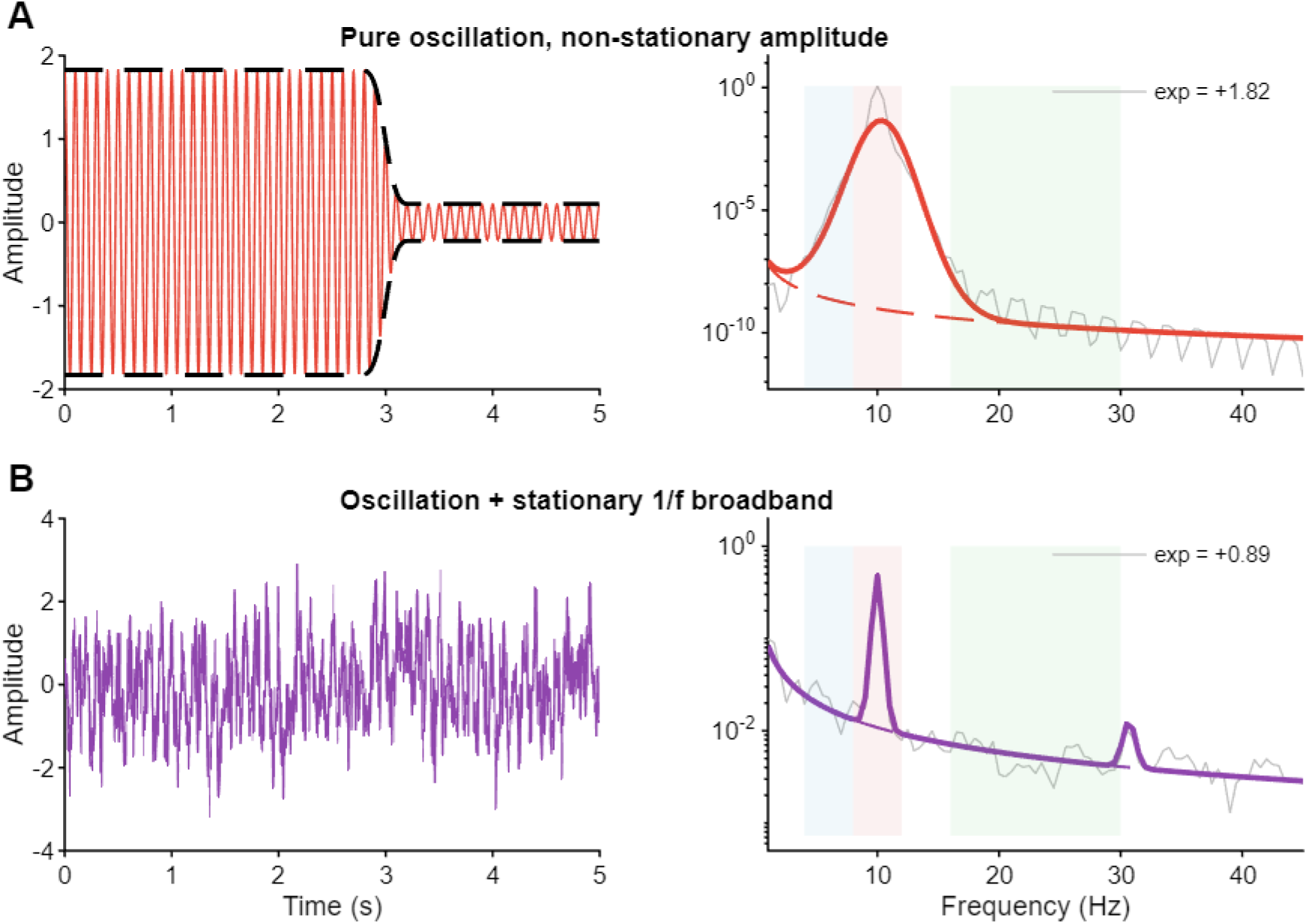
The spectral non-identifiability problem. Two signals shown in 5 s time-domain traces (left) with their power spectral densities (right, light grey) and specparam fixed-mode fits (coloured lines, dashed = aperiodic component). Shaded regions: theta (blue), alpha (red), beta (green). (A) Pure alpha oscillation with non-stationary amplitude (ERD-like piecewise suppression, Gaussian-smoothed); no aperiodic component in the generating process. (B) Stationary alpha oscillation superimposed on a 1/f-like broadband process (NeuroDSP sim_powerlaw input exponent −1, i.e. a specparam-convention exponent of +1; the two toolboxes use opposite sign conventions). The two spectra differ quantitatively, the broadband signal returns a higher specparam exponent than the AM signal, but they share the same qualitative form (peak plus sloping baseline), and no feature of the spectrum identifies which generative process produced each. The decomposition assigns an aperiodic exponent to (A) despite the absence of any aperiodic component in the signal.

Figure 1 is a minimal illustration of a more general property. A continuum of generative processes (stationary broadband activity, transient or amplitude-modulated oscillations, episodic bursts, mixtures of these) will all be summarised by specparam as a peak (or several) plus an aperiodic exponent. The processes differ in their temporal structure, but the spectrum collapses that temporal information into a small set of frequency-domain statistics that do not, in general, identify which process produced the signal. For panel A, the positive exponent arises geometrically: a non-stationary amplitude envelope translates into sidebands around the carrier, and the line that specparam fits in log-log coordinates across the off-peak region has a positive slope. The exact magnitude of this slope is not a fixed property of the amplitude modulation; it depends on the envelope shape and on what other broadband content fills the fitting range. The point of Figure 1 is not that AM is always a major source of inflated exponents in practice, but that the spectrum, summarised as peak plus exponent, does not contain enough information to identify the temporal structure that produced it.

### 3.2 Features extracted from a signal

Figure 2 illustrates the features extracted from a signal using one illustrative scenario (bursty beta on a 1/f background, scenario E). Panel A shows the raw signal with clear burst episodes. Panel B shows the power spectrum, the slope (+0.94) and beta peak are consistent with either bursting or genuine broadband activity. Panel C shows the beta amplitude envelope with high CV = 0.54, reflecting the burst on/off structure. Panel D shows fBOSC burst abundance concentrated at 21 Hz (0.38) with low values elsewhere. Panel E shows the SPRiNT exponent timecourse with SD = 0.09. The same set of features is applied to all scenarios and empirical case studies that follow.

**Figure 2.**
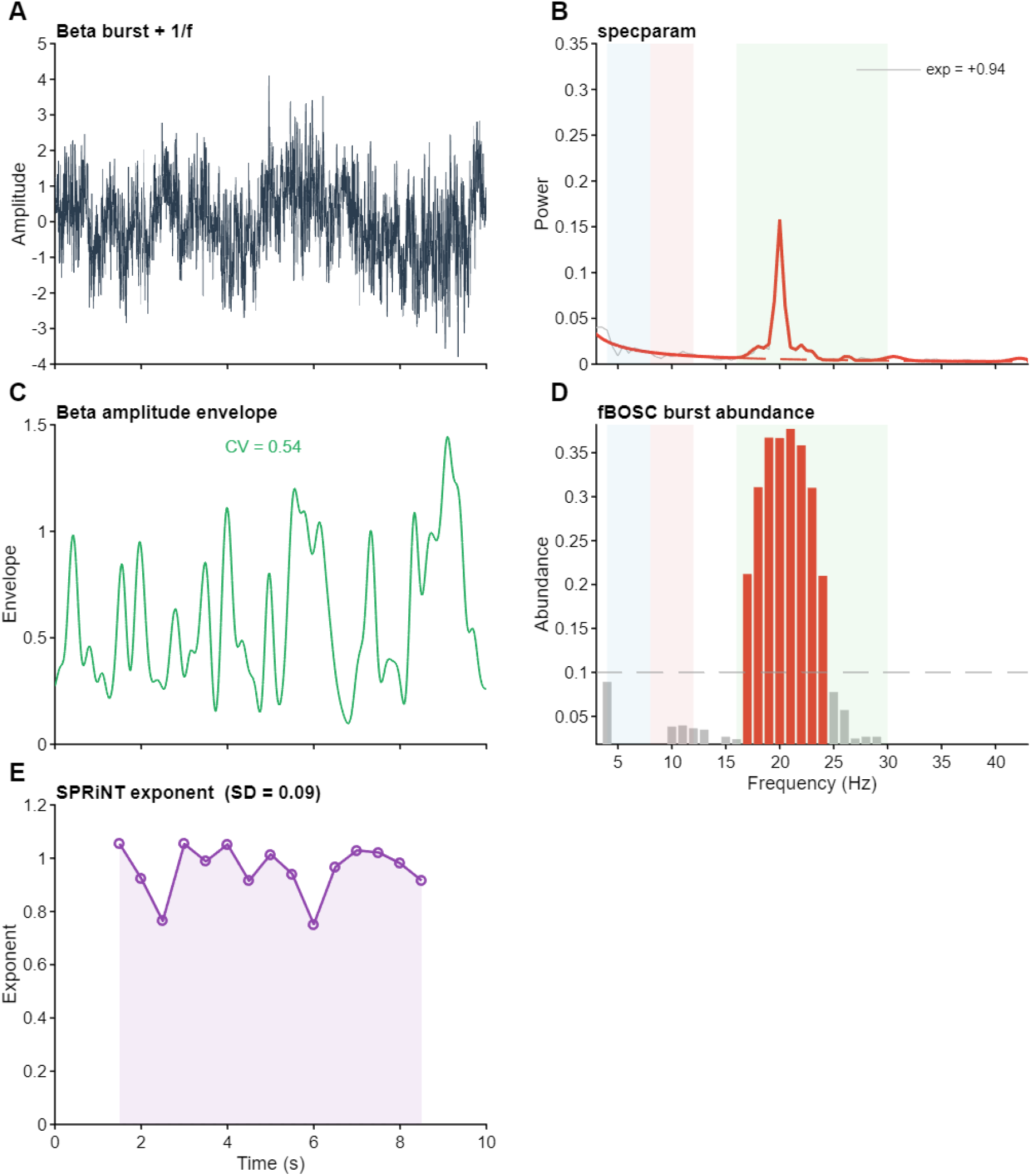
Feature extraction applied to scenario E (beta burst + 1/f-like background). (A) Raw 10 s signal. (B) PSD with specparam fit (exp = +0.94, R² = 0.95). (C) Beta-band Hilbert amplitude envelope; CV = 0.54, reflecting burst on/off dynamics. (D) fBOSC burst abundance per frequency; elevated at 21 Hz (0.38), low elsewhere. (E) SPRiNT aperiodic exponent over time (SD = 0.09).

### 3.3 What a 1/f-like spectrum can look like in the time domain

The same family of spectral statistics - a peak plus a sloping baseline - is consistent with several distinct temporal structures. Table 1 shows feature values for the five illustrative scenarios.

**Table 1.** Feature values for the five illustrative scenarios. Exponent: specparam fixed-mode aperiodic exponent. CV (α) and CV (β): Hilbert envelope coefficient of variation in alpha (8-12 Hz) and beta (16-30 Hz). SPRiNT SD: standard deviation of the SPRiNT aperiodic exponent across 1 s STFT windows. fBOSC peak: peak burst abundance and frequency. R² is shown as “—” for scenarios A and B because the spectrum is near-flat at exp ≈ 0, where the fixed-mode aperiodic fit is uninformative and its R² is not meaningfully interpretable; it is reported for the scenarios with a genuine slope (C–E).

| Scenario | exp | R<sup>2</sup> | CV ($\alpha$) | CV ($\beta$) | SPRiNT SD | fBOSC peak |
| --- | --- | --- | --- | --- | --- | --- |
| **A. Pure $\alpha$ oscillation** | -0.02 | — | 0.09 | — | 0.09 | — |
| **B. AM $\alpha$ oscillation** | -0.05 | — | 0.70 | — | 0.08 | — |
| **C. $\alpha$ on 1/f background** | +0.76 | 0.95 | 0.19 | — | 0.11 | — |
| **D. Bursty $\beta$** | +0.21 | 0.85 | — | 0.85 | — | — |
| **E. Bursty $\beta$ on 1/f background** | +0.94 | 0.95 | — | 0.54 | 0.09 | 0.38 @ 21 Hz |

Each scenario produces a different combination of features. Scenario A (pure α): exponent is essentially zero (slightly negative; a flat noise floor with a single peak yields an exponent near or just below zero) and the envelope is flat. Scenario B (AM α): exponent is still near zero (the sinusoidal modulator creates sidebands that specparam absorbs as part of the peak, not as a slope), but the envelope CV jumps to 0.70 - the AM is visible in the envelope, not in the spectrum. Scenario C (α + 1/f): the exponent climbs to +0.76 driven by the broadband component; the envelope CV stays low because the oscillation itself is stationary. Scenario D (bursty β): the exponent is moderately positive (+0.21) and the beta envelope CV is high (0.85), capturing the on/off structure. Scenario E (bursty β + 1/f): exponent is +0.94, envelope CV in beta is 0.54 (lower than scenario D because the 1/f background fills the off-burst intervals), and fBOSC shows concentrated burst abundance at 21 Hz.

No single feature fully characterises the temporal structure underlying a 1/f-like spectrum. The exponent is positive in scenarios C and E and moderately positive in D, but the temporal structures differ: C has a stationary oscillation on a broadband background, D has episodic bursts without one, E has both. The envelope CV is elevated in B, D, and E, distinguishing them from the stationary scenarios A and C, but does not tell you whether the elevated CV reflects sinusoidal AM, irregular bursting, or the combination. fBOSC localises burst activity in frequency but is uninformative when the broadband baseline is near zero, as in A and B. Together the features constrain the space of plausible interpretations more than any one alone: the practical recommendation is to report several, precisely because different processes leave different signatures across them, and the combination narrows the interpretive possibilities in ways that no single summary statistic can.

### 3.4 Case study: alpha lateralisation

Figure 3 shows specparam applied to a trial-averaged spectrogram of the simulated alpha lateralisation paradigm (Melcón et al. 2025), for the contralateral (60% amplitude suppression, ERD) and ipsilateral (60% enhancement, ERS) dipoles; a constant-amplitude control was also computed and is reported in the statistics but not plotted. Panel A: trial-averaged signals, with the amplitude modulation visible in the ERD/ERS window. Panel B: the aperiodic exponent rises for the contra ERD (baseline 1.40 → ERD window 2.05, Δ = +0.65) and barely changes for the ipsi ERS (1.43 → 1.36, Δ = −0.07); the constant-amplitude control does not move (1.41 → 1.41).

**Figure 3.**
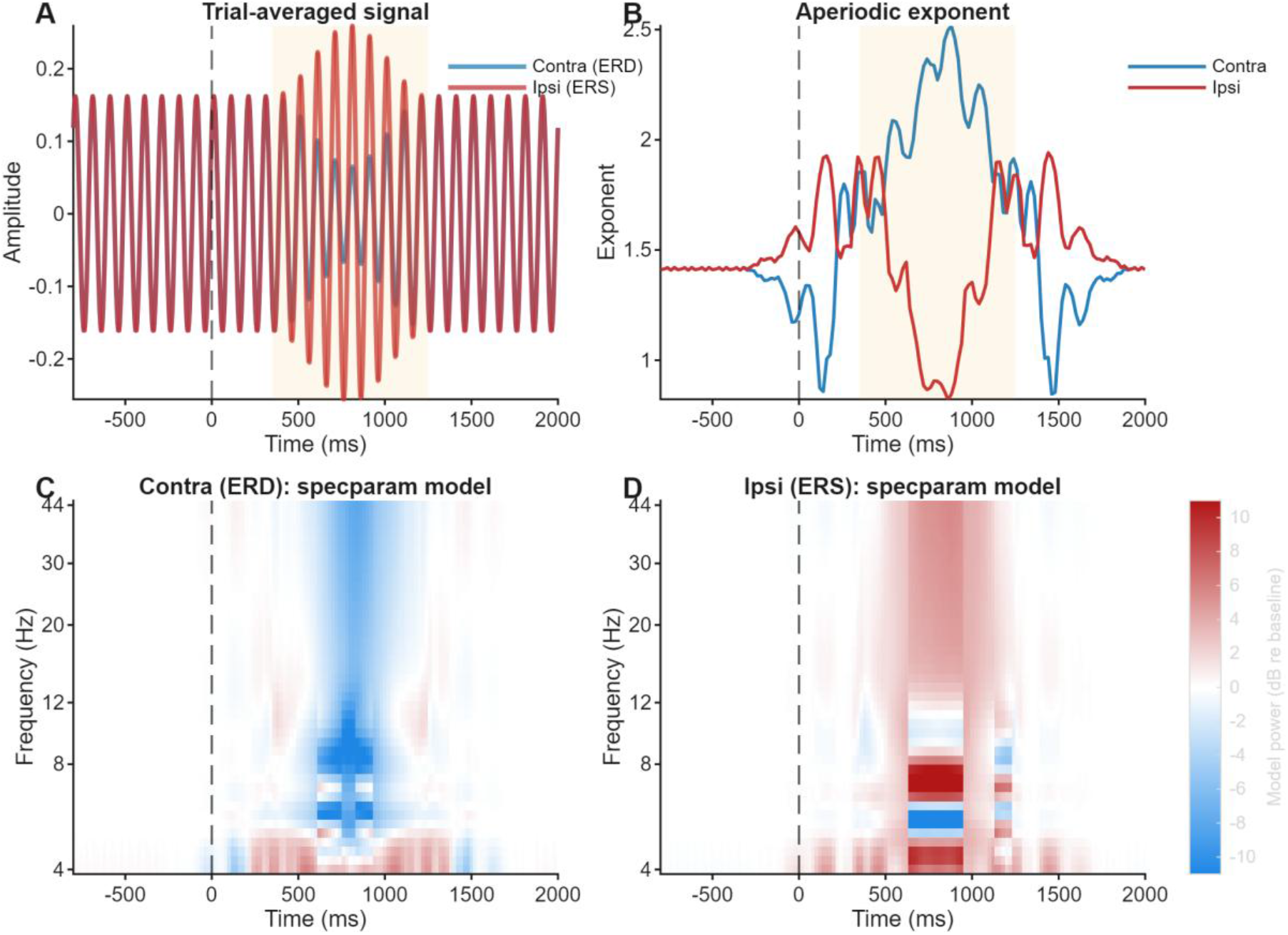
Trial-averaged spectral parameterisation of the simulated alpha lateralisation paradigm (60 trials, 10 Hz carrier, no noise, no frequency jitter, carrier phase drawn per trial and shared across conditions; signals generated on a 2 s-padded time base and cropped to the −800 to 2000 ms display window after the wavelet transform; ERD/ERS depth 60%, Hann-shaped over 350–1250 ms). Contra = ERD (blue), ipsi = ERS (red). (A) Trial-averaged signal. (B) Aperiodic exponent (sampled every 10 ms; effective resolution ∼225 ms at 10 Hz): contra 1.40 → 2.05 (Δ +0.65); ipsi 1.43 → 1.36 (Δ −0.07). Mean R² = 0.98–0.99 across unmasked bins. (C, D) specparam-modelled spectrum (the fitted model, not the raw wavelet power) for the contra and ipsi conditions, in dB relative to the pre-stimulus baseline; shared colour scale, white = edge-masked bins. The modelled change is confined to the modulation window. Under the ipsi ERS the fitted peak power falls (peak-window Δ = −0.43 log units) despite the oscillation growing, and the fitted peak centre frequency drifts from 9.7 to 12.6 Hz (Δ = +2.9 Hz); under the contra ERD the fitted peak power falls (Δ = −0.98) with the centre near 9.3–9.7 Hz. Orange shading in A/B marks the ERD/ERS window.

Mean R² = 0.98–0.99 across unmasked bins. Panels C and D show the specparam-modelled spectrum (the fitted model, not the raw wavelet power) for the contra and ipsi conditions, expressed in dB relative to the pre-stimulus baseline. Because the amplitude modulation is confined to the 350–1250 ms Hann window, the modelled change is restricted to that interval, giving the panels their block-like time structure; within the window the change reverses sign across frequency, because an exponent increase pivots the modelled power law about a crossover frequency (raising low-frequency and lowering high-frequency power relative to baseline) rather than shifting it uniformly.

This connection between oscillatory amplitude modulation and the trial-averaged waveform is the basis of the baseline-shift account of evoked responses (Nikulin et al., 2007, 2010): if an oscillation has a non-zero mean, then any modulation of its amplitude shifts the mean of the signal, and this shift survives trial averaging as a slow deflection, an evoked response, whereas a zero-mean oscillation would cancel. Oscillations and (slow) ERPs are, on this view, two sides of the same process rather than separate generators.

The exponent change here is not a change in some underlying broadband state; it is a consequence of how specparam fits the spectrum when oscillatory power changes relative to the broadband floor. The behaviour of the fitted alpha peak makes this concrete and is itself a cautionary finding. The fitted peak power does not track the true oscillatory amplitude: for the contra ERD it falls (peak-window Δ = −0.98 log units, as expected when the oscillation is suppressed), but for the ipsi ERS it also falls (Δ = −0.43) even though the oscillation grows. The reason is visible in the fitted centre frequency: under the strong ERS the fit reorganizes from two peaks near 9.7 Hz to a single, broader peak whose centre drifts to about 12.6 Hz (Δ = +2.9 Hz), while for the contra and baseline conditions it stays near 9.7 Hz. The corrected oscillatory power and centre frequency returned by specparam therefore cannot be read as faithful measures of oscillatory amplitude when the amplitude change is large, because the peak model itself is unstable in that regime (see 4.3). The constant-amplitude control, by contrast, shows no movement in either the exponent or the fitted peak, so the changes in the ERD/ERS conditions are driven by the amplitude modulation and not by the analysis.

### 3.5 Case study: auditory oddball ERP

Figure 4 shows the trial-averaged parameterisation applied to ECoG recordings from a sensor on the auditory cortex of a marmoset during a roving oddball protocol (Komatsu et al. 2015). The N1 complex peaked at −147.2 µV at 59 ms. The grand average PSD yielded a steep specparam exponent (+2.61, R² = 0.96, 2 peaks), reflecting broadband low-frequency power of the ERP transient, with a dominant peak at 22.5 Hz. Because the grand average spans only 0.35 s, this is a single-window, low-resolution PSD (frequency spacing ∼2.9 Hz over [1, 45] Hz), so the exponent and peak frequency are coarse descriptors rather than precise estimates; the substantive result is the time-resolved exponent below. The trial-averaged aperiodic exponent changed only modestly across the unmasked epoch (baseline 1.22, N1 window 1.36, Δ = +0.13 from pre-stimulus baseline to the N1 window), a much smaller change than the contra ERD in the lateralisation simulation (Δ = +0.65).

**Figure 4.**
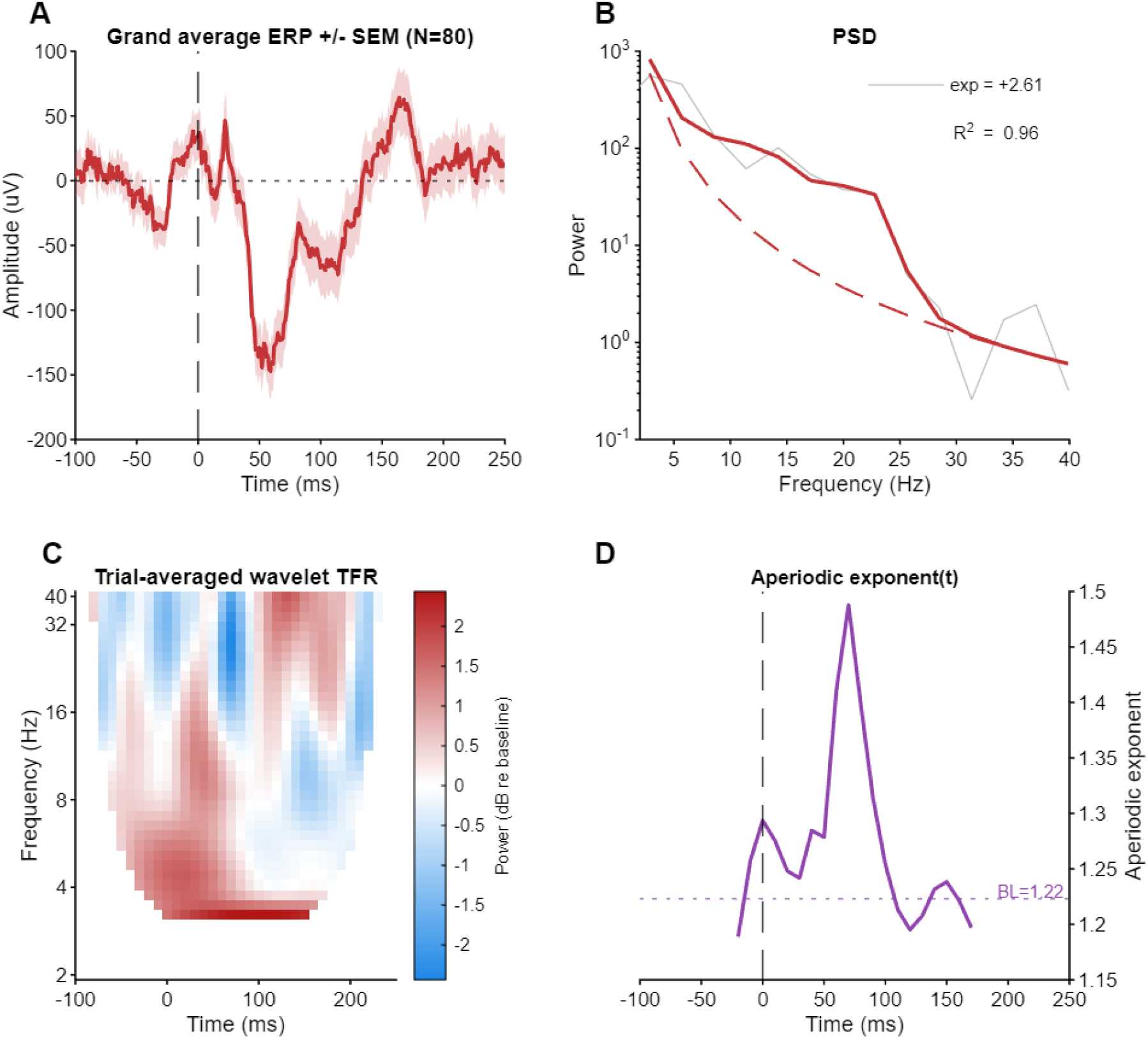
Empirical roving oddball ERP (256 Hz, 80 deviant trials, auditory cortex, marmoset; Komatsu et al. 2015). (A) Grand average +/- SEM; clear N1 and P2 components visible; N1 = −147.2 µV at 59 ms. (B) Grand average PSD with specparam fixed fit (exp = +2.61, R² = 0.96, 2 peaks); dominant peak at 22.5 Hz. (C) Trial-averaged spectrogram with frequency-dependent edge masking (white = NaN within FWHM/2 of epoch boundary). (D) Aperiodic exponent timecourse (unmasked range −20 to 170 ms); baseline 1.22, N1 window 1.36, Δ = +0.13; mean R² = 0.99.

The auditory N1 is a broadband transient that elevates power across many frequencies approximately proportionally, preserving the spectral slope. The alpha ERD is narrowband: it selectively reduces power at 10 Hz while the broadband floor is unchanged, altering the slope. In this single-channel example the trial-averaged parameterisation thus narrows the interpretive possibilities, separating a selective (oscillatory) from a broadband (transient) account of the power change within a single epoch; yet, as with the lateralisation simulation, it constrains rather than determines the interpretation.

### 3.6 Case study: scalp EEG vs ECoG, rest vs sedation

Results obtained from simultaneous scalp EEG and ECoG resting recordings from the same posterior temporal location (scalp T5 and an adjacent ECoG channel; Oosugi et al., 2017), are reported in Figure 5 in two states: resting wakefulness and propofol sedation, 300 s each. Unlike the trial-averaged wavelet parameterisation used in Sections 3.4–3.5, the exponents reported here are obtained from Welch power spectra and from SPRiNT applied to the continuous recordings; the absolute values are therefore those of the standard estimators and are not subject to the constant-Q wavelet frequency-response offset that affects the trial-averaged wavelet exponents in Sections 3.4–3.5 (see §4.3).

**Figure 5.**
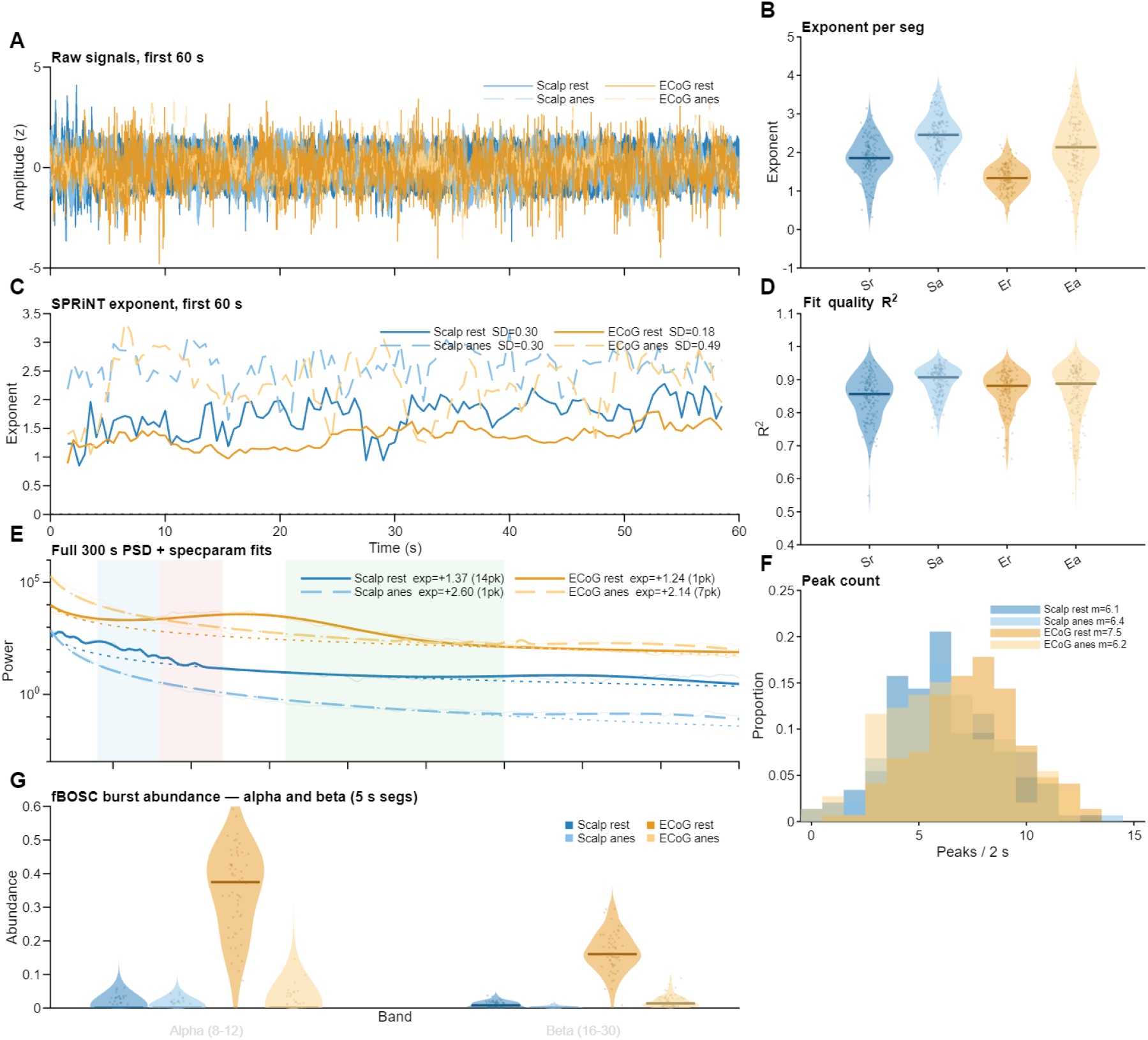
Simultaneous scalp EEG and ECoG (256 Hz, 300 s) under resting wakefulness and propofol sedation. Blue = scalp (solid: rest, dashed: sedation); orange = ECoG (solid: rest, dashed: sedation). (A) Raw signals, first 60 s, z-scored. (B) Aperiodic exponent distribution across 2 s segments (Sr = scalp rest, Sa = scalp sedation, Er = ECoG rest, Ea = ECoG sedation). (C) SPRiNT aperiodic exponent, first 60 s; SD values in legend. (D) specparam R² per 2 s segment. (E) Full 300 s PSD with specparam fits (solid) and aperiodic components (dotted); exponent and peak count in legend. (F) Peak count per 2 s segment; means in legend. (G) fBOSC burst abundance across 5 s segments for alpha (8–12 Hz) and beta (16–30 Hz); distributions shown as violins across segments.

The full 300 s power spectra (Panel E) show a large exponent increase under sedation in both modalities: scalp rest exp = +1.37 (14 peaks), scalp sedation exp = +2.60 (1 peak), ECoG rest exp = +1.24 (1 peak), ECoG sedation exp = +2.14 (7 peaks). The segment-level exponent distributions (Panel B, 2 s segments) show that this increase is consistent across the recording: scalp rest mean = 1.84 ± 0.53, scalp sedation mean = 2.48 ± 0.49, ECoG rest mean = 1.34 ± 0.34, ECoG sedation mean = 2.11 ± 0.74. In this recording ECoG sedation shows the highest variability, plausibly reflecting genuine non-stationarity of the anaesthetic state. A coupling between the high-frequency spectral slope and alpha-band amplitude, including under propofol, has been reported directly in monkey ECoG by Muthukumaraswamy & Liley (2018).

The time-resolved exponent (SPRiNT, Panel C) shows a different picture across modalities. ECoG rest has the lowest variability (SD = 0.18), ECoG sedation the highest (SD = 0.49), a pattern consistent with the state contrast visible in Panel B. Scalp rest and scalp sedation have indistinguishable SPRiNT SD (0.30 in both, to two decimals; this could be coincidental at n = 1), suggesting that at scalp the variability is dominated by an artefact floor that is independent of neural state, consistent with the observations of Ameen et al. (2025) on iEEG versus scalp EEG.

The fBOSC analysis (Panel G, 5 s segments, alpha and beta bands) shows a clear modality-by-state interaction. At rest, ECoG has detectable alpha bursts (median abundance across segments 0.384, IQR 0.286–0.478) and beta bursts (median 0.103, IQR 0.060–0.145). Under sedation both collapse to near zero (alpha median 0.000, beta median 0.000, IQR up to 0.029). At scalp, abundance is near zero in all conditions and does not differentiate rest from sedation in this recording.

Read together, the panels illustrate a dissociation that is central to the paper’s argument. The spectral slope increases substantially under sedation in both modalities, with the same direction, and a similar magnitude. But the temporal correlates of that change differ: at ECoG the burst structure collapses, the SPRiNT variability increases, and the segment-level exponent distributions shift in a way that is consistent with a change in the nature of the broadband activity. At scalp none of these temporal correlates are detectable, and the SPRiNT trace is indistinguishable across states. The slope change is evident in both cases. For ECoG, the temporal features narrow the interpretive possibilities: the burst collapse and SPRiNT variability increase are consistent with a shift in broadband activity rather than a pure AM change. For scalp, the burst structure that would make the contrast visible is absent, making the temporal features less informative.

## 4. Discussion

### 4.1 Behaviour versus mechanism

The analyses in this paper describe the observable temporal structure of the signal at the measurement scale; they are not claims about underlying neural mechanisms. The power spectrum, and any finite set of time-series statistics, cannot uniquely identify the generative process: multiple mechanisms produce indistinguishable observable signatures. This is the distinction drawn by the mechanism-vs-behaviour framework (Rosas et al., 2022; Marinazzo, 2025): the observable behaviour of a system can be characterised without thereby identifying the mechanism that produces it. What can be characterised is whether the oscillatory amplitude shows temporal patterns consistent with modulation or bursting, or whether it is consistent with a stationary broadband process. These are descriptions that constrain the space of plausible mechanisms without selecting among them.

The value of characterising the observable behaviour at this finer level lies in its practical consequences. Specparam extracts a periodic component by subtracting an aperiodic baseline; if the baseline is inflated by AM-induced broadband spread, the extracted peaks are biased, regardless of mechanism. SPRiNT tracks the exponent over time; if the exponent fluctuates in a way correlated with oscillatory amplitude dynamics, treating the exponent as a readout of some underlying broadband state is behaviourally unjustified. The practical correction is the same: examine whether the broadband behaviour is predominantly AM-driven, and if so, treat the oscillatory amplitude timecourse rather than the exponent as the primary variable of interest.

The question "why bother with this distinction if you cannot tell the difference anyway?" has a direct answer: even when the distinction is difficult, knowing its difficulty is itself informative. A finding that the envelope of a detected oscillation is highly variable in the conditions where the exponent also changes is grounds for caution. A finding that the envelope is stable while the exponent moves is grounds for treating the exponent change more seriously as a property of the broadband background. The features do not deliver mechanism, but they constrain interpretation.

### 4.2 The role of stationarity and scale

Whether a process appears stationary depends on the observation window. A sinusoidal AM process with a 10 s modulation period will appear essentially stationary in a 2 s window but clearly non-stationary in a 30 s window. The illustrative scenarios in Section 3 use 10 s epochs, which captures AM at the 1 Hz modulation frequency (10 cycles) but would not resolve AM at 0.1 Hz. In practice, the relevant modulation timescale is set by the cognitive or neural process under investigation.

This scale-dependence bounds what the envelope CV can detect: it measures the variability of the Hilbert envelope at timescales from a few cycles of the carrier frequency up to the recording window. At the fast end of that range, burst activity with very short (200 ms) bursts at irregular intervals will not produce a smooth envelope modulation; fBOSC is better suited for detecting this than the CV feature, which requires sustained, regular amplitude variations.

The extreme of the timescale argument is the spike: a single action potential has a 1/f-like spectrum (Gerstein & Mandelbrot, 1964; Bédard et al., 2006). When many cells fire independently, their summed spectrum also approximates 1/f. When cells are synchronised, their summed activity shows oscillatory peaks. The spectrum is thus a function of both the single-cell dynamics and the degree of synchrony, and it changes across timescales. The analyses presented here operate at the mesoscopic scale of LFP/EEG/ECoG and do not resolve the cell-level contributions. This is a fundamental scope limitation, not a methodological failure.

### 4.3 Trial-averaged parameterisation: implications and limitations

The trial-averaged parameterisation reduces estimator variance by averaging across trials rather than smoothing across time, which leaves the wavelet-determined resolution unchanged. With N = 60 trials, the per-bin power-estimate variance is reduced by a factor of 60 (standard error by √60 ≈ 7.7×), sufficient for reliable specparam fits (mean R² = 0.98–0.99 across unmasked bins in the lateralisation simulation; Section 3.4). The timecourse is sampled every 10 ms, but the effective resolution remains the wavelet FWHM (∼225 ms at 10 Hz). Within the short-epoch domain the relevant comparison is with per-trial windowed parameterisation (Section 2.4); SPRiNT and STPPTO address a different regime, tracking exponent drift and burst thresholds in long continuous recordings.

Because the lateralisation simulation is noise-free and jitter-free, the absolute exponent (∼1.40 at baseline) is the constant-Q wavelet’s frequency response, not aperiodic content of the signal; accordingly, only within-condition relative changes from the trial-averaged wavelet parameterisation are interpretable, and no absolute exponent from this method should be read as a measure of broadband activity. The exponent rise during the contra ERD in the alpha lateralisation simulation (Δ = +0.65) is a new observable. It reflects the fact that when alpha power falls relative to the broadband background (ERD), the local spectral slope between the peak and the background steepens; conversely, when alpha power rises relative to the background, the slope appears shallower. This is why the amplitude-increasing ipsi ERS does not inflate the exponent (Δ = −0.07). This is a measurement property, not necessarily a neural state change, and should be interpreted with caution. The contrast with the auditory N1 (exponent change small, Δ = +0.13) is instructive: the N1 is a broadband transient that elevates power across many frequencies simultaneously, preserving the relative slope; alpha ERD is narrowband and reshapes the slope.

Some technical aspects and potential limitations are worth mentioning: frequency-dependent temporal smearing (FWHM ≈ 562 ms at 4 Hz) makes it unsuitable for theta-band events shorter than ∼500 ms; the wavelet constant-Q bandwidth introduces frequency-dependent peak amplitude distortion relative to Welch PSDs, but this is systematic and consistent across conditions. A specific caution follows from the lateralisation simulation (Figure 3): when the oscillatory amplitude change is large, specparam’s peak model becomes unstable, so the fitted peak parameters do not track the true oscillation. Under the ipsi ERS the fitted peak power falls even though the oscillation grows, and the fitted peak centre frequency drifts upward (here from about 9.7 to 12.6 Hz) as the fit reorganises from two narrow peaks to a single broad one; the aperiodic component itself barely moves over the same window. Corrected oscillatory power and peak frequency should therefore not be read as faithful amplitude or frequency measures in this regime. The two single-channel case studies presented here illustrate the approach but do not establish its generality across participants, paradigms, or recording sites. The apparent onset timing of a detected modulation will be shifted backward by approximately FWHM/2 relative to the true onset, as a consequence of wavelet temporal smearing. This is not a bias in the statistical sense; it is the fundamental Gabor uncertainty of time-frequency analysis.

### 4.4 A possible reframing of studies of aperiodic activity

A substantial literature frames research questions as investigating the role of aperiodic activity in phenomenon X, or reports that aperiodic activity is correlated with cognitive variable Y, typically introduced with the observation that while specific frequency bands have an established functional role, the contribution of aperiodic activity has not been studied in detail. This framing, while useful for motivating specparam-based analyses, conflates the two levels of description that the mechanism-vs-behaviour framework (Section 4.1) keeps distinct. When a study correlates the specparam exponent with a cognitive variable, it is describing a relationship between two observable behaviours, not a relationship between a mechanism and a function. This is a legitimate and informative description, but it should not be interpreted as establishing that some specific neural mechanism (E/I balance, membrane dynamics, firing rate) is functionally engaged, unless independent evidence for the mechanism is provided.

Three practical considerations follow. First, the aperiodic component returned by specparam is a feature of the power spectrum, not a directly observable neural process. Its value depends on the frequency range over which the model is fitted, the success of peak detection, the choice of fixed vs knee model, and any preprocessing that modifies the spectral slope. None of these dependencies are properties of the neural signal; they are properties of the analysis pipeline. A study reporting a correlation between the aperiodic exponent and a cognitive variable should establish that the finding is robust to reasonable variations in these parameters, and ideally report both fixed and knee fits.

Second, the analytical-choice problem identified by da Silva Castanheira, Landry & Fleming (2025) deserves emphasis. Their practical recommendation is to use specparam-modelled rhythmic power rather than detrended band power whenever rhythmic and arrhythmic components are to be related independently. This recommendation is followed in the present paper: the alpha-power values reported in Section 3.4 (e.g. modelled alpha power in contra and ipsi conditions) are specparam-modelled Gaussian peak amplitudes, not detrended band powers. Studies that report band-by-band changes derived from log- or linear-detrending should be re-examined with this concern in mind.

Third, a finding that "the aperiodic exponent changes between conditions" could mean that the broadband spectral slope has genuinely changed, or it could mean that an oscillatory peak has changed in amplitude or bandwidth in a way that was not fully captured by specparam’s periodic model. The case studies in Section 3 illustrate exactly this ambiguity. The alpha lateralisation simulation shows an exponent increase driven by a narrowband power decrease (the contra ERD); the auditory N1 shows a broadband transient with only a small exponent change (Δ = +0.13). The two are distinguishable from the temporal envelope of the carrier band and from the time-resolved aperiodic exponent. Studies reporting aperiodic exponent changes should routinely include the amplitude envelope CV and the time-resolved (SPRiNT) exponent stability as corroborating evidence, and A concrete instance is the proposal of Preston, Smith and Voytek (2026) that task-evoked decreases in trial-to-trial neural variability reflect decreases in the aperiodic exponent (a flattening of the spectrum). The same observable is also what a non-stationary oscillatory envelope produces: a stimulus-locked amplitude change alters both trial-to-trial variability and the fitted exponent without requiring a change in a separate aperiodic process. The two readings are not distinguishable from the variability decrease or the exponent shift alone, and separating them again requires the temporal structure of the envelope.

Burst detection methods (see (Power et al., 2026) for a review) represent a special case of this general problem. All amplitude-based burst detection approaches require an accurate estimate of the broadband 1/f-like background before setting a burst detection threshold. The temporal-structure features used throughout this paper address the prior question that these methods presuppose answered: is the broadband background a genuinely non-oscillatory process, or is it itself a consequence of the amplitude non-stationarity of the oscillations being detected? If the latter, the specparam background estimate is inflated by the very signal it is meant to exclude.

### 4.5 The aperiodic/periodic division and band-frequency conventions

The blurring of the periodic/aperiodic boundary is symmetric in both directions. Non-stationary oscillatory amplitude leaks into the broadband slope; broadband 1/f activity, filtered into any narrow band, returns a signal whose amplitude varies over time, resembling a genuinely modulated oscillation. The coefficient of variation of the envelope alone does not separate the two; however, the full envelope distribution can in principle distinguish them, since narrowband-filtered Gaussian noise has a Rayleigh-distributed envelope with a fixed coefficient of variation of √((4−π)/π) ≈ 0.523 (Rice, 1944), whereas deterministic sinusoidal AM produces a different envelope distribution. These classical results on narrowband random processes are the deeper reason the two are hard to separate: any narrowband random process can be written as a slowly-varying amplitude and phase modulating a carrier, so a stretch of filtered broadband noise is itself indistinguishable, sample by sample, from an amplitude- and phase-modulated oscillation. “Broadband” and “oscillatory” are, at this level, two descriptions of the same narrowband signal rather than two generators. Reporting the envelope distribution or its shape, not only its CV, is therefore a constructive strengthening. The two phenomena are not independent confounds but mirror images of the same underlying fact: the temporal envelope and the spectral slope are coupled, and changing one necessarily affects the other. This is why no single feature can identify the mechanism. The features used in this paper work by triangulating across descriptors that respond differently to the two directions of mixing: envelope CV is high in both AM-modulated oscillations and bandpass-filtered 1/f, but specparam fit quality, peak detectability, and SPRiNT exponent variability distinguish them.

This symmetry is in tension with the way the two components are typically reported in the empirical literature. Most studies treat the oscillatory peak parameters and the aperiodic parameters as separate predictors that can each be related to behaviour, cognition or clinical status (e.g. Donoghue et al., 2020; Schaworonkow, 2023; da Silva Castanheira et al., 2025; Marsicano et al., 2026). Even studies that explicitly model both components and find that they jointly account for the same outcome generally conclude, on the basis of mediation analyses or partial-correlation arguments, that each contributes "independent" variance and therefore reflects a distinct underlying mechanism. The position taken here is that this reading goes beyond what the data support: if a non-stationary oscillatory amplitude generates a broadband slope, and a broadband process generates an envelope that looks AM, then a residual independent contribution of the "aperiodic" component, after the oscillatory component is partialled out, does not unambiguously index a separate physiological mechanism, because the partition itself depends on a decomposition that the signal does not in general support. The empirical counterpart to this position is van Engen et al. (2026), who parameterise single-trial working-memory spectra and argue that event-related changes in aperiodic activity can masquerade as changes in band-limited oscillatory power, in particular that the mixed theta findings in the literature partly reflect aperiodic dynamics. Their conclusion runs in the same direction as the review discussed in Section 4.5 (oscillatory effects reattributed to aperiodic activity) and rests on the same premise this paper questions: that the two are separable components the signal licenses one to disentangle. On the reading taken here their result is better stated as underdetermination, since the spectrum they fit is equally consistent with an aperiodic change and with amplitude modulation of the oscillation, and their two-window design, like any static-window parameterisation, cannot access the temporal signature that would even constrain which is which.

A parallel blurring occurs between oscillations and evoked responses. Under the baseline-shift mechanism, late, slow ERP components arise from amplitude modulation of non-zero-mean oscillations rather than from an independent additive response (Nikulin et al., 2007, 2010; Mazaheri & Jensen, 2008). The same amplitude dynamics that reshape the spectral slope therefore also generate part of the evoked response, reinforcing the view that the conventional partitions (periodic/aperiodic, oscillation/ERP) cut across a single underlying set of amplitude dynamics.

The band-frequency framework adds a further discretisation: it assigns frequencies to labelled categories (theta, alpha, beta, gamma) with fixed boundaries, which are then treated as functionally distinct entities. This is epistemically convenient but biologically arbitrary. A change in alpha-band power that is not accompanied by a specparam-detected alpha peak may reflect a shift in the aperiodic exponent rather than a change in alpha oscillatory amplitude. Band-by-band analyses that use raw bandpass power rather than specparam peak amplitude are susceptible to this misinterpretation.

### 4.6 Limitations

The empirical examples in this paper are deliberately limited in scope. The auditory oddball ERP (one channel, one participant, one paradigm), the alpha lateralisation simulation (a model, not data), and the simultaneous scalp-ECoG comparison (one channel pair, one participant, two states) illustrate how different temporal structures project into the same family of spectral statistics. They are not, and should not be read as, general characterisations of any of those modalities. Every empirical statement in Sections 3.4–3.6 should be read as conditional on the specific recording analysed.

The scenarios in Section 3.3 are illustrative. They are not meant to be a calibration set, a validation set, or a model of any specific neural process. The temporal structures they instantiate (sustained sinusoid, sinusoidal AM, sinusoid on 1/f, beta burst, beta burst on 1/f) are caricatures chosen to make a conceptual point about the diversity of time-domain configurations consistent with a similar spectral shape. Real neural signals will fall on a continuum between these caricatures and combinations of them.

The framework assumes approximately sinusoidal oscillations: non-sinusoidal waveforms produce harmonics that complicate envelope estimation and may mimic broadband activity in some frequency ranges (Schaworonkow, 2023). The trial-averaged parameterisation is limited by frequency-dependent temporal smearing. fBOSC abundance is uninformative when the aperiodic background is near zero. No biophysical model is provided linking the observable features to specific circuit mechanisms; the §4.7 discussion describes what such a model would have to address.

### 4.7 Biophysical context

The question of what generates neural 1/f spectra has a long history that predates specparam. Pascual-Marqui et al. (1988) proposed the Xi-Alpha model, decomposing the EEG spectrum into two additive independent processes: Xi (the broadband background, always present) and Alpha

(the oscillatory peak, not always present. Crucially, they provided a partial generative interpretation of the Xi process: its spatial coherence decays exponentially with interelectrode distance and is real-valued (zero phase), consistent with a homogeneous, isotropic stochastic process driven by short-range cortical interactions.

Wang et al. (2026) extend the Xi-Alpha decomposition to the bispectrum, generalising the oscillatory (peak) term and denoting it Rho, using BiSpectral EEG Component Analysis (BiSCA), jointly fitting the power spectrum and bispectrum. Their central finding is that the Xi (aperiodic) component behaves as a linear, Gaussian stochastic process at the quadratic level, while the Rho (oscillatory) components are the primary source of nonlinear resonance. This provides bispectral evidence for a distinction that the present paper approaches from the time domain: when a broadband spectral slope is genuinely not driven by oscillatory AM, it should behave as a stationary, linear process, characterized by low envelope CV, stable SPRiNT exponent, and negligible bicoherence.

A further implication of this lineage is that the spectral trend may not be a power law at all. Pascual-Marqui et al. (1988) already parametrised the background not as 1/f but with an explicit closed form that decays smoothly with a characteristic corner frequency rather than as a straight line in log-log coordinates; Wang et al. (2026) generalise this to a Cauchy/Student-t kernel, and Brake et al. (2024) independently derive the unitary spectrum as a sum of decaying Lorentzian terms set by receptor kinetics. On all of these accounts a straight line in log-log space is a local approximation to a curved, decaying spectrum rather than evidence of a scale-free process, and the knee parameter that specparam introduces is essentially a partial re-introduction of the corner frequency that the pure power-law form discards. This matters because "1/f" is not a neutral description: it carries connotations of self-similarity, criticality, and fractal scaling that a finite-corner, decaying spectrum does not. Reading a fitted exponent as a power-law exponent therefore risks attributing scale-free dynamics to a process whose generative model is an ordinary linear, finite-timescale stochastic system, consistent with (Wang et al. 2026)’s The same additive, finite-timescale picture underlies the corrected decomposition of Bloniasz et al. (2026): because the periodogram of a Gaussian process is Gamma-distributed and its components add in linear power, they fit a summed broadband-plus-Gaussian-peak model with a Gamma likelihood rather than a Gaussian fit to log-power, and recover broadband and rhythmic parameters that specparam mis-assigns when rhythms are strong (software: SL_specdecomp). Their broadband term is itself a knee/Lorentzian-type decay rather than a pure power law, consistent with the lineage above.

A complementary source of spectral complexity is the contribution of non-sinusoidal waveform shape to apparent higher-frequency oscillations. Schaworonkow (2023) demonstrates that a substantial proportion of beta-band spectral peaks in human EEG are not genuine beta oscillations but harmonics of non-sinusoidal alpha rhythms: in the eyes-closed condition, 65.6% of participants show a beta-peak frequency that is twice the alpha-peak frequency. The bispectrum provides a more direct tool for detecting this: harmonic coupling between a fundamental at f and its second harmonic at 2f appears as a peak in the bicoherence at the frequency pair (f, f), which BiSCA (Wang et al., 2026) uses via joint spectrum-bispectrum fitting.

A biophysical forward-modelling approach is taken by Brake et al. (2024), who address directly whether the EEG spectral trend can arise from genuinely arrhythmic neural activity. Their core finding is that the unitary EEG spectrum follows a sum of two Lorentzian functions, with the two timescales governed by the deactivation kinetics of GABA and AMPA receptors. Crucially, they show that scalp EEG cannot reflect truly asynchronous neural activity: dipoles from individual neurons average to zero unless the underlying synaptic currents are at least partially correlated. The broadband EEG trend therefore arises not from uncorrelated spike trains or self-organised criticality, but from correlated synaptic currents whose timescales shape the spectral envelope. This was validated empirically using propofol, a GABA receptor agonist that slows inhibitory synaptic decay and shifts the spectral trend in a quantitatively predictable way. They further show that shifts in synaptic properties can corrupt apparent oscillatory power estimates, providing a concrete mechanism by which a change in the broadband background can masquerade as a change in a narrowband rhythm.

The pedagogical review of Milotti (2002) shows that there is no single canonical mechanism behind 1/f spectra: superposition of exponential relaxation processes with rates distributed over a finite interval, diffusion with appropriate boundary conditions, self-organised criticality, and certain deterministic dynamical systems all yield approximately 1/f scaling. The simplest of these is also the most directly relevant here: if a process is built from many exponential relaxations whose rates are uniformly distributed in some interval, the resulting spectrum is approximately 1/f within that range. This shifts the question from "what universal mechanism produces 1/f" to "what is the distribution of timescales in the system being measured, and is it broad enough to look 1/f over the observable band."

Evertz et al. (2022) apply exactly this superposition-of-relaxations logic to resting EEG, with a neurally motivated twist: instead of overdamped exponential relaxations they use damped alpha-band linear oscillators driven by independent stochastic forcing. By solving an inverse problem on the EEG power spectrum, they recover the distribution of damping rates needed to reproduce the observed shape, and find that this distribution is typically bimodal. The weakly-damped mode produces the narrowband alpha peak; the heavily-damped mode, where oscillators relax rapidly and contribute broad, low-Q resonances, produces the 1/fβ tail. Eyes-closed to eyes-open alpha blocking is then accounted for by an increase in the damping of the weakly-damped mode, rather than by any separate change in a distinct 1/f process. In this picture both features emerge from a single family of damped oscillatory processes, and a change in the "aperiodic" tail need not signal an independent physiological process. The same damped-oscillator account was advanced earlier by Muthukumaraswamy & Liley (2018), who additionally showed that the high-frequency spectral slope co-varies with alpha-band power over time in MEG, EEG, and ECoG, and shifts under propofol and other pharmacological agents, a direct empirical demonstration that the broadband slope and oscillatory amplitude are dynamically coupled rather than independent.

These considerations converge on a deeper epistemic point. Brake et al. (2024) present the most mechanistically explicit case for the "two radically different things" view: the broadband EEG trend is generated by correlated synaptic currents with specific receptor-kinetic timescales, a process categorically distinct from the synchronous oscillatory dynamics that generate narrowband peaks. Evertz et al. (2022), by contrast, show that both peak and slope can be accounted for by a single population of damped oscillators whose damping distribution spans from underdamped to overdamped. At the single-neuron and single-synapse level, the same spike train, depending on the degree of synchrony across the population, the timescale of observation, and the spatial summation by volume conduction, can contribute to either the oscillatory peak or the broadband slope, or to both simultaneously. There is no a priori reason why the same microscale mechanism should produce only one or the other at the macroscale. Whether the 1/f slope and the oscillatory peaks are "two different things" is not a question that can be answered from a power spectrum, or from the temporal structure of the signal, without additional bottom-up knowledge of the generative architecture. Without that knowledge, which is typically unavailable in scalp EEG recordings, the most honest statement is that the power spectrum is consistent with either account, and that the temporal structure of the signal constrains, but does not uniquely determine, which is dominant at the measurement scale.

## 5. Conclusions

The spectral slope of neural recordings cannot be attributed to a specific generative mechanism from time-series analysis alone. The temporal structure of the signal, represented by envelope behaviour, exponent stability, burst abundance, can be characterised, and that characterisation constrains how a spectral slope change should be interpreted, without delivering a mechanism. Multiple temporal structures map to the same family of spectral statistics, as the five illustrative scenarios in Section 3.3 show, and the three case studies show that this ambiguity is not removed by moving to real data: in some recordings the temporal features are informative; in others they are not. Applying specparam to a trial-averaged spectrogram extends the time-resolved approach to short event-related epochs, where the exponent timecourse can register the difference between a selective oscillatory amplitude change and a broadband transient—a contrast not available in the static spectrum, though, as the lateralisation simulation shows, these exponent dynamics are a measurement property of the analysis rather than a direct readout of the underlying processes.

The practical message for empirical research is conservative: when reporting a change in the "aperiodic" component, examine the temporal envelope of any detected oscillations in the same data, compare fixed and knee specparam fits where a knee is plausible, prefer modelled peak power to detrended band power (while recognising, as the lateralisation simulation shows, that even modelled peak power becomes unreliable when the amplitude change is large enough to destabilise the peak fit; the two cautions address different failure modes), and treat the discrete labels of "periodic" and "aperiodic" as conventions for summarising a continuous signal rather than as separate physiological entities. The framework presented here does not deliver a classifier, and is not intended to. It identifies what to look at and what to be cautious about.

The periodic/aperiodic framing has been productive precisely because it provided a tractable decomposition of an otherwise featureless spectral slope. What it does not provide is a partition of neural processes: the same spectrum is consistent with a stationary broadband background, with amplitude-modulated oscillations, with bursts, and with mixtures of these. The instability cuts deeper still: the “aperiodic” side of the partition need not even follow a power law— generative models of the trend (Pascual-Marqui et al., 1988; Brake et al., 2024; Wang et al., 2026) describe it as a decaying spectrum with a characteristic corner rather than a scale-free 1/f process (a finite-timescale view shared even by proponents of distinct aperiodic activity: Preston, Smith and Voytek (2026) likewise reject a self-organized-criticality reading of neural 1/f on the grounds that neural spectra carry a characteristic knee rather than unbroken power-law scaling), so the dichotomy fixes neither the temporal nature of the oscillatory side nor the spectral form of the trend. A recent review by Preston, Smith and Voytek (2026) sets out the opposing position in its strongest form, treating periodic and aperiodic activity as distinct processes with separate physiological generators, and arguing that much of the field has conflated them. Notably, it frames the conflation in the opposite direction to the present paper: that effects previously attributed to oscillations may be better explained by aperiodic activity.

The concern here is the converse: changes attributed to a distinct aperiodic process can, at the level of the observable signal, be re-expressions of non-stationary oscillatory amplitude. That the conflation can be argued in either direction from the same spectra is consistent with the view taken here: if an exponent change is equally compatible with a shift in a broadband generator and with amplitude modulation of an oscillation, the partition is not settled by the spectrum, and the question is one for the time domain. A related critique has been made at the level of the estimator itself: Bloniasz et al. (2026) note that field potentials arise from the additive (linear) superposition of biophysical generators, so their power spectra sum, whereas specparam combines a broadband term and oscillatory peaks multiplicatively in log-power space, implicitly assuming that one process scales another. Under strong rhythms this misspecification biases both the estimated peak and the estimated trend, and can attenuate or distort genuine changes in the trend; the same observation that motivates my time-domain argument (broadband and oscillatory power are not independently readable from the spectrum) appears there as a bias in the spectral decomposition itself. The static/dynamic distinction operates on the temporal structure of the signal rather than on its spectrum, and it is this structure that determines how a spectral slope change should be interpreted: whether as a shift in broadband activity, a change in oscillatory amplitude dynamics, or an artefact of analysis. The framework presented here is one way to make that assessment; its principal message is that the assessment is necessary.

## Data and code availability

All analysis code is available at https://github.com/danielemarinazzo/aperiodic_AM. The empirical datasets are openly available via NeuroTycho (https://neurotycho.org/, Nagasaka et al., 2011).

